# Deglycosylation of isoflavone *C*–glucoside puerarin by combination of two recombinant bacterial enzymes and 3–oxo–glucose

**DOI:** 10.1101/816074

**Authors:** Kenichi Nakamura, Shu Zhu, Katsuko Komatsu, Masao Hattori, Makoto Iwashima

## Abstract

*C*–Glucosides are resistant to glycoside hydrolase activity because the anomeric carbon of glucose is directly connected to aglycone via carbon-carbon bonding. A human intestinal bacterium strain PUE related to *Dorea* species can metabolize the isoflavone *C*–glucoside puerarin (daidzein 8–*C*–glucoside) to daidzein and glucose by more than three bacterial enzymes which have not been well-characterized. We previously reported that 3”–oxo–puerarin is an essential reaction intermediate in enzymatic puerarin degradation and characterized a bacterial enzyme of DgpB–C complex which cleaved the *C*–glycosidic bond in 3”–oxo–puerarin. However, the exact enzyme catalyzing the oxidation of C–3” hydroxyl in puerarin has not been identified, and the other metabolite corresponding to the precursor of D–glucose, derived from the sugar moiety in 3”–oxo–puerarin in the cleaving reaction catalyzed by the DgpB–C complex, remains unknown.

In this study, we demonstrated that recombinant DgpA, a Gfo/Idh/MocA family oxidoreductase, catalyzed puerarin oxidation in the presence of 3–oxo–glucose as the hydride accepter. In addition, enzymatic *C*–deglycosylation of puerarin was achieved by a combination of recombinant DgpA, DgpB–C complex, and 3–oxo–glucose. Furthermore, the metabolite derived from the sugar moiety in 3”–oxo–puerarin cleaving reaction catalyzed by DgpB–C complex was characterized as 1,5–anhydro–D–*erythro* –hex–1–en–3–ulose, suggesting that the *C*–glycosidic linkage is cleaved through a *β*–elimination like mechanism.

**Importance:** One important role of the gut microbiota is to metabolize dietary nutrients and supplements such as flavonoid glycosides. Ingested glycosides are metabolized by intestinal bacteria to more absorbable aglycones and further degradation products which show beneficial effects in humans. Although numerous glycoside hydrolases that catalyze *O*–deglycosylation have been reported, enzymes responsible for *C*–deglycosylation are still limited. In this study, we characterized enzymes involved in *C*–deglycosylation of puerarin from a human intestinal bacterium PUE. To our knowledge, this is the first report of the expression, purification and characterization of an oxidoreductase involved in *C*–glucoside degradation. This study provides new insights for the elucidation of mechanisms of enzymatic *C*–deglycosylation.

## Introduction

More than 1000 species of bacteria colonize the human gut and affect host health and diseases (1, 2). One important role of the gut microbiota is to metabolize dietary nutrients and supplements such as flavonoids (3, 4). Many natural flavonoids in plants are stored in the form of glycosides. In general, ingested glycosides are poorly absorbed in the human small intestine because of their hydrophilicity but are reported to be metabolized by intestinal bacteria to more absorbable aglycones and further degradation products (3, 4). For example, the isoflavone *O*–glucoside daidzin (daidzein 7–*O*–glucoside) is hydrolyzed to aglycone and the resulting daidzein is reduced to (*S*)–equol, which shows beneficial effects in humans by preventing hormone-related diseases (4–6). From this perspective, naturally occurring glycosides are considered to be a type of prodrug activated by intestinal bacterial metabolism (7).

*C*–Glucoside is a naturally occurring glycoside in which the anomeric carbon of glucose is directly connected to the aglycone via carbon-carbon bonding. Because of the stability of *C*–glucosyl bonds, *C*–glucosides are resistant to glycoside hydrolase and acid treatments, in contrast to *O*–glucosides. Although the catalytic mechanisms of enzymatic *C*–deglycosylation have not been well-characterized, some intestinal bacteria were reported to metabolize *C*–glucosides to the corresponding aglycones (8–14). Braune *et al*. reported that heterologous expression of five *Eubacterium cellulosolvens* genes (*dfgABCDE*) in *Escherichia coli* led to metabolization of flavone *C*–glucosides to aglycone (15). This was the first study in which the genes involved in *C*–deglycosylation were cloned; however, the roles of these 5 gene products in the reaction remain unclear.

We previously isolated a human intestinal bacterium PUE (92% similarity in 16S rRNA gene sequence with *Dorea longicatena)*, which metabolizes the isoflavone *C*–glucoside puerarin (daidzein 8–*C*–glucoside) to daidzein and glucose (10, 16). Enzymatic studies revealed that more than three bacterial enzymes involved in multi-step reaction of *C*–deglycosylation (17). Moreover, a putative puerarin-metabolizing-operon composed of 8 genes (*dgpA*–*H*) from strain PUE was identified (accession number LC422372), and recombinant DgpB–C complex was shown to cleave the *C*–glycosidic bond in 3”–oxo–puerarin but not puerarin (Fig. 1) (18). These results indicated that 3”–oxo–puerarin is an essential reaction intermediate in puerarin degradation reaction, and an unidentified oxidoreductase that catalyzes oxidation at the C–3” hydroxyl of puerarin was predicted to be encoded in the operon.

**Fig. 1.**
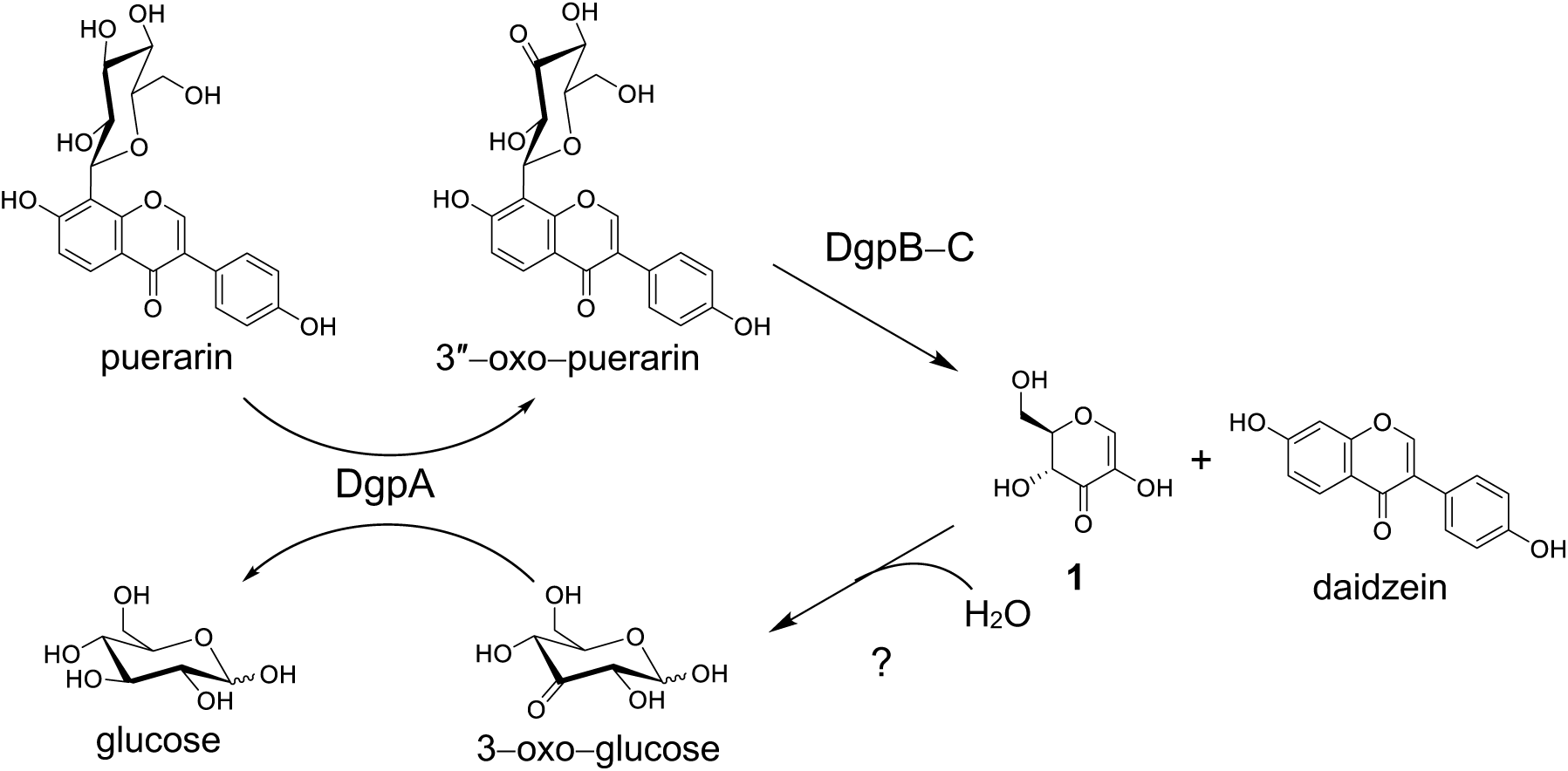
Proposed puerarin degradation pathway catalyzed by DgpA and DgpB–C complex.

In this study, we demonstrated that recombinant DgpA catalyzed puerarin oxidation in the presence of 3–oxo–glucose as the hydride accepter (Fig. 1). In addition, enzymatic *C*–deglycosylation of puerarin was achieved by a combination of DgpA, DgpB–C complex and 3–oxo–glucose. Furthermore, the real metabolite derived from the sugar moiety in 3”–oxo–puerarin catalyzed by DgpB–C complex was characterized as 1,5–anhydro–D–*erythro* –hex–1–en–3–ulose (**1**).

## Materials and methods

### Chemicals and materials

Puerarin was purchased from Carbosynth Limited. 1,2:5,6–di–*O*–isopropylidene–*α*–D–glucofuranose was obtained from TCI. NAD^+^, NADH, NADP^+^, and NADPH were purchased from Oriental Yeast Co., Ltd. 3”–oxo–puerarin was prepared as previously described (18). Genomic DNA of strain PUE was obtained according to the literature (18). Recombinant DgpB–C complex was prepared as previously reported (18).

### Preparation of 3–oxo–glucose

3–oxo–glucose was synthesized according to the literature procedures (19, 20). Briefly, C–3 hydroxyl of 1,2:5,6–di–*O*–isopropylidene–*α*–D–glucofuranose was oxidized using NaOCl, nor–AZADO and KBr in CH_2_Cl_2_/aq NaHCO_3_ (19). The obtained 1,2:5,6–di–*O*–isopropylidene–3–oxo–α–D–glucofuranose was treated with trifluoroacetic acid:H_2_O (9:1) to give 3–oxo–glucose (20).

### Construction of recombinant DgpA expression vector

A DNA fragment encoding *dgpA* gene was amplified from genomic DNA of strain PUE by PCR using forward primer (5’–AAA<u>GAATTC</u>ATGAGTAAATTAAAAATTGG–3’, EcoR I site is underlined) and reverse primer (5’–AAA<u>CTCGAG</u>TTAGAATTTAATTGTCTCAT–3’, Xho I site is underlined). The amplified fragment was cloned into the EcoR I/ Xho I site of the pET–21a (+) vector. A nucleotide sequence encoding N–terminus T7 tag of the constructed vector was removed by deletion PCR using forward primer (5’–TATACATATGAGTAAATTAAAAATT–3’) and reverse primer (5’– TTACTCATATGTATATCTCCTTCTTA –3’) according to the manufacturer’s instructions of a PrimeSTAR Mutagenesis Basal kit (Takara Bio Inc.).

### Expression and purification of recombinant DgpA

The constructed vector was transformed into *E. coli* BL21 (DE3) and the transformant was cultured at 37°C in LB broth containing 100 μg/mL ampicillin. A recombinant DgpA was induced with 1 mM isopropyl β–D–thiogalactopyranoside and the culture was continued at 25°C for 15 h. The cells were disrupted by sonication and centrifuged to obtain a supernatant containing crude recombinant DgpA.

### Purification of recombinant DgpA

Recombinant DgpA was purified by two-step column chromatography of an anion exchange column chromatography (HiPrep Q FF 16/10 column, GE Healthcare) and a hydrophobic column chromatography (HiPrep Butyl FF 16/10 column, GE Healthcare). The purified protein was dialyzed against 50 mM potassium phosphate buffer (pH 7.4).

### Measurement of UV-vis absorption spectrum of the purified DgpA

UV**-**vis absorption spectrum of the purified DgpA (0.5 mg/mL in 50 mM potassium phosphate buffer, pH 7.4) was recorded using a spectrophotometer UV-1800 (Shimadzu, Japan).

### Determination of DgpA-bound NAD(H)

Determination of DgpA-bound NAD(H) was performed according to the literature procedure with minor modification (21). To the 1 mg of purified DgpA in 0.1 mL 50 mM potassium phosphate buffer (pH 7.4) was added 0.9 mL methanol, which was stored at 0°C for 15 min. The solution was passed through 0.22 μm membrane and the filtrate was concentrated in vacuo to approximately 0.1 mL to remove methanol. To the concentrated solution was added H_2_O (0.1 mL) and passed through 0.22 μm membrane. The filtrate containing dissociated NAD(H) from DgpA was analyzed by high-performance liquid chromatography (HPLC). HPLC conditions were as follows: column, COSMOSIL 5C_18_–MS–II (Nacalai Tesque) 4.6×150 mm; flow rate, 1 mL/min; detection, 260 nm; mobile phase, (A) 20 mM sodium dihydrogen phosphate and (B) acetonitrile (linear gradient from 0% to 10% B concentration over 30 min); injection volume, 10 μL.

### Enzyme assay

A reaction mixture (100 μL) consisting of an enzyme (DgpA with or without DgpB–C complex, 1 μg each), a substrate (puerarin or 3”–oxo–puerarin, 0.5 mM), and an additive (glucose or 3–oxo–glucose, 5 mM) in 50 mM potassium phosphate buffer (pH 7.4) was incubated at 37°C for 30 min. Methanol (300 μL) was added to the reaction solution and metabolites were analyzed by ODS–HPLC. HPLC conditions were the same as previously described (18).

### Purification and structure determination of a reductive metabolite of 3”–oxo–puerarin catalyzed by DgpA

A reaction mixture (10 mL) including DgpA (100 μg), 3”–oxo–puerarin (1 mM), and glucose (50 mM) in 50 mM potassium phosphate buffer (pH 7.4) was incubated at 37°C for 60 min. The reaction solution was passed through Amicon Ultra–15 10K centrifugal filter devices (Merck Millipore Ltd.) and the obtained low molecular fraction was acidified with 1 mol/L HCl. The acidified solution was applied to inertSep C18 column (GL Sciences) and washed with H_2_O, and then eluted with methanol. The methanol fraction was concentrated in vacuo to give a reductive metabolite. ^1^H and ^13^C NMR spectra were identical to that of puerarin standard.

### Structure determination of a metabolite derived from the sugar moiety of 3”–oxo–puerarin catalyzed by DgpB–C complex

A reaction mixture (30 mL) containing DgpB–C complex (1.8 mg) and 3”–oxo–puerarin (18.6 mg) in H_2_O was incubated at 37°C for 30 min. The resulting precipitate (daidzein) was removed by filtration. To the filtrate was added 10 mL of water saturated butan–1–ol (containing 0.1% AcOH) and then liquid–liquid partition was carried out. The water layer was concentrated to approximately 3 mL and applied to inertSep C18 column eluting with H_2_O. The eluent was concentrated in vacuo to give 1,5–anhydro–D–*erythro*–hex–1–en–3–ulose (**1**, 1.7 mg).

^1^H and ^13^C nuclear magnetic resonance (NMR) spectra were recorded with Varian NMR system 600 and the residual solvent of CD_3_CN was used as an internal standard (^1^H, 1.93 ppm; ^13^C, 1.3 ppm). ^1^H NMR of **1** (600 MHz, CD_3_CN) δ: 3.75 (1H, dd, *J*=4.3, 12.7 Hz, one of H-6), 3.83 (1H, dd, *J*=2.1, 12.7 Hz, another one of H-6), 4.01 (1H, dddd, *J*=0.5, 2.2, 4.3, 13.3 Hz, H-5), 4.33 (1H, d, *J*=13.3 Hz, H-4), 7.36 (1H, s, H-1). ^13^C NMR of **1** (150 MHz, CD_3_CN) δ: 61.4 (C-6), 68.6 (C-4), 84.5 (C-5), 135.2 (C-2), 148.0 (C-1), 191.5 (C-3).

## Results

### Expression and purification of DgpA, Gfo/Idh/MocA family oxidoreductase

3”–oxo–puerarin is a key intermediate in the enzymatic *C*–deglycosylation of puerarin (Fig. 1); however, the exact enzyme catalyzing the oxidation of C–3” hydroxyl in puerarin has not been identified. We previously reported the putative puerarin-metabolizing-operon composed of 8 genes (*dgpA*–*H*) from intestinal bacterium strain PUE (18). DgpA (BBG22493.1) and DgpF (BBG22498.1), both regarded as gene products of the operon suggested as closely related to oxidoreductase in the Gfo (glucose–fructose oxidoreductase) / Idh (inositol 2–dehydrogenase) / MocA (rhizopine catabolism protein MocA) protein family (22). Particularly, DgpA was implicated in puerarin oxidation because the *dgpA* gene deduced amino acid sequence at the N-terminus was identical to that of a previously reported protein involved in puerarin metabolism (17).

To characterize the enzymatic activity of DgpA, the encoding gene *dgpA* was heterologously expressed in *E. coli*, and the recombinant protein was purified by two-step column chromatography. In sodium dodecyl sulfate-polyacrylamide gel electrophoresis (SDS-PAGE) analysis, the purified DgpA appeared as a single band with an apparent molecular mass of 42 kDa, showing good agreement with the calculated molecular mass of 40,161 Da (Fig. 2, lane 1). The recombinant DgpB–C complex which catalyzes the deglycosylation of 3”–oxo–puerarin was also analyzed by SDS–PAGE (Fig. 2, lane 2).

**Fig. 2.**
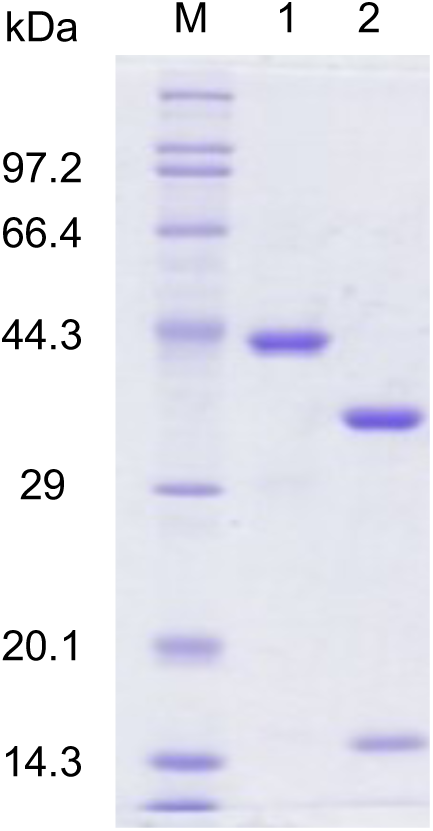
SDS-PAGE analysis of recombinant proteins used in this study. Lane M, marker proteins; lane 1, recombinant DgpA (40,161 Da); lane 2, recombinant DgpB–C complex (DgpB: 16,047 Da, DgpC: 36,883 Da). SDS-PAGE was carried out with 12.5% polyacrylamide gel.

### Determination of DgpA-bound NAD(H)

In the UV-vis spectrum of purified DgpA, a broad shoulder peak at approximately 340 nm was observed, suggesting that nicotinamide cofactors such as NAD(H) or NADP(H) were bound to the enzyme (Fig. 3). To characterize the cofactors, HPLC analysis was performed after the protein was treated with cold methanol to dissociate the cofactors. As shown in Fig. 4b, two major peaks were observed at 10.9 and 13.5 min in HPLC analysis of DgpA**-**bound cofactors. These two peaks were characterized as NAD^+^ and NADH by comparing the retention times to authentic nicotinamide cofactors (Fig. 4a). These results indicate that NAD(H) functioned as the cofactor which binds tightly but non-covalently to DgpA.

**Fig. 3.**
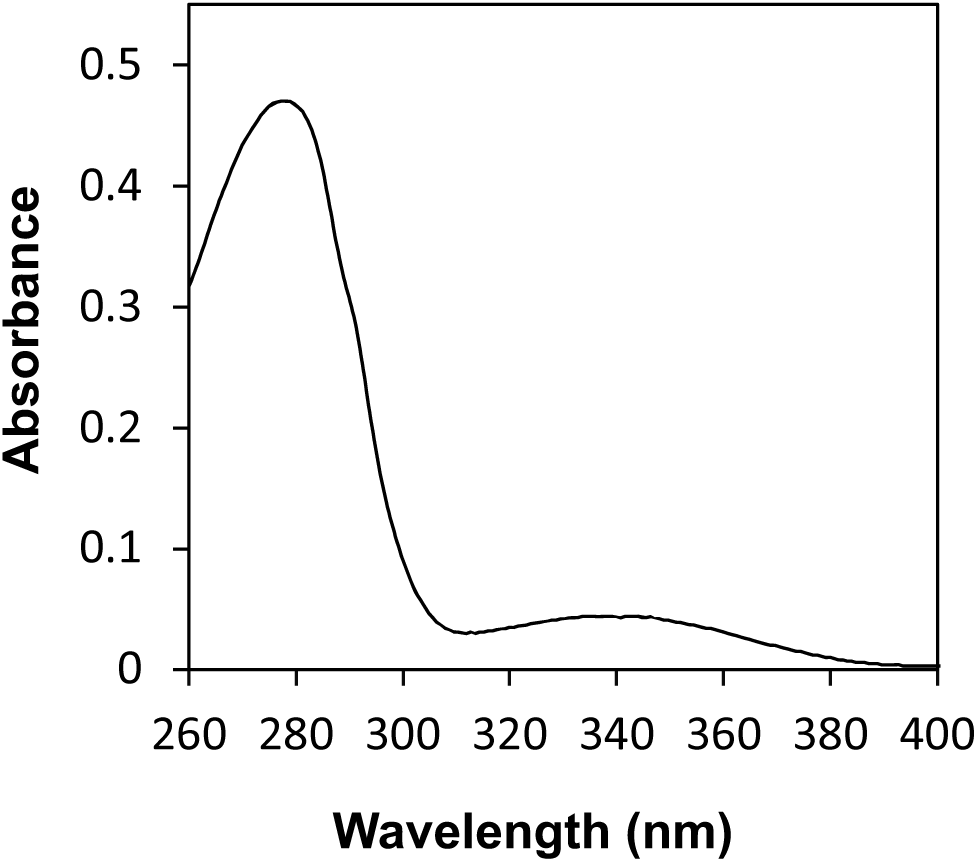
UV-vis absorption spectrum of the purified DgpA.

**Fig. 4.**
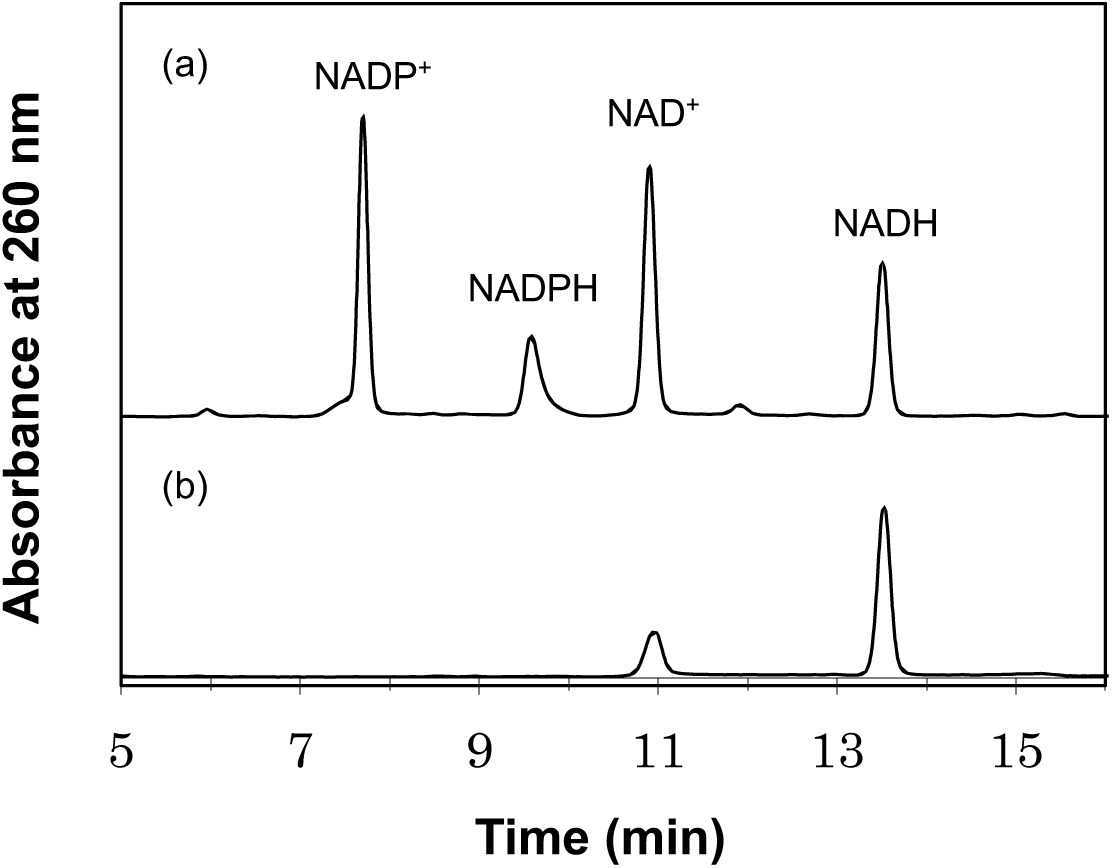
HPLC analysis of DgpA**-**bound NAD(H). Purified DgpA was denatured with cold methanol and the dissociated cofactors were analyzed by HPLC. (a) Authentic mixture of cofactors, NAD^+^, NADH, NADP^+^, and NADPH, (b) DgpA**-**447 bound cofactors.

### Oxidation of puerarin by DgpA and 3–oxo–glucose

To confirm the oxidation of puerain catalyzed by DgpA, recombinant DgpA was incubated with 0.5 mM puerarin in potassium phosphate buffer (pH 7.4) at 37°C for 30 min. According to HPLC analysis, no metabolites were detected under this condition (Fig. 5a). The same result was obtained when 1 mM NAD^+^ and 1 mM MnCl_2_ were added to the reaction mixture, despite these two cofactors have been reported to increase the enzymatic activity during puerarin *C*–deglycosylation (17). As shown in Fig. 1, DgpA may require 3–oxo–glucose for oxidation of puerarin, as the ultimate sugar metabolite in puerarin degradation should be glucose rather than an oxo-sugar derivative (16). Based on this assumption, 3–oxo–glucose was added to the reaction mixture including DgpA and puerarin, resulting in the detection of two metabolite peaks at 8.1 and 8.6 min in HPLC analysis (Fig. 5b). The retention times and elution profiles of the metabolites were identical to those of 3”–oxo–puerarin in the buffer, which easily isomerized to a mixture of the 3”–oxo form (a peak at 8.1 min), 2”–oxo form, and its intramolecular-cyclic acetal (a peak at 8.6 min, overlapping) as previously reported (18). These results demonstrate that DgpA catalyzed oxidation at the 3”–hydroxyl of puerarin by using 3–oxo–glucose as the hydride accepter (Fig. 1).

**Fig. 5.**
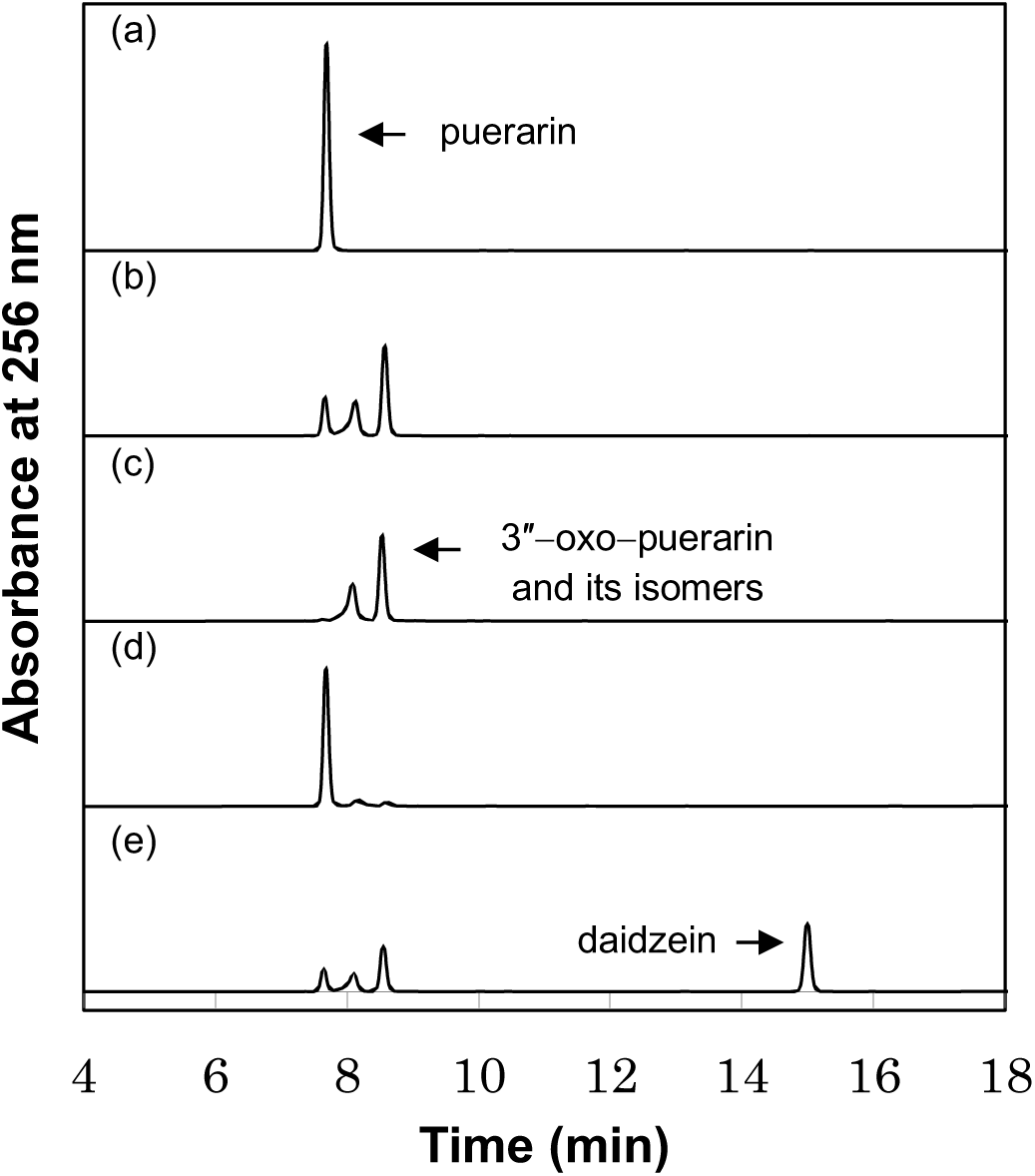
HPLC analysis of enzymatic reaction metabolites catalyzed by DgpA and DgpB–C complex. Enzymatic reaction mixtures were incubated at 37°C for 30 min and the reaction solution were analyzed by HPLC. The composition of the reaction mixtures were as follows, (a) puerarin and DgpA, (b) puerarin, DgpA, and 3–oxo–glucose, (c) 3″–oxo–puerarin and DgpA, (d) 3″–oxo–puerarin, DgpA, and glucose, (e) puerarin, DgpA, DgpB–C complex, and 3–oxo–glucose.

### Reduction of 3”–oxo–puerarin by DgpA and glucose

To identify the actual metabolites in the oxidation reaction by DgpA, an enzymatic counterreaction was proposed. 3”–Oxo–puerarin standard and DgpA were incubated with or without D–glucose at 37°C for 30 min, followed by HPLC analysis (Fig. 5c, d). In the reaction of 3”–oxo–puerarin standard and DgpA with glucose, one conspicuous metabolite peak was detected at 7.7 min by HPLC (Fig. 5d). After chromatographic isolation, the metabolite structure was confirmed as puerarin, but not 3”–axial–hydroxyl epimer (D–allose type *C*–glycoside), based on NMR analysis. These findings indicate that the reaction catalyzed by DgpA was reversible and the metabolites in the puerarin oxidation reaction were verified as 3”–oxo–puerarin and D–glucose, as shown in Fig. 1.

### *C*–Deglycosylation of puerarin by a combination of DgpA, DgpB–C complex, and 3–oxo–glucose

DgpB–C complex was reported to metabolize 3”–oxo–puerarin to daidzein (18). To achieve enzymatic *C*–deglycosylation of puerarin, two recombinant bacterial enzymes (DgpA and DgpB–C complex) and 3–oxo–glucose were incubated with puerarin at 37°C for 30 min, which was analyzed by HPLC. The peak detected as daidzein was observed at 15.0 min in the HPLC chromatogram (Fig. 5e), indicating that *C*–deglycosylation of puerarin was accomplished by the recombinant enzymes.

### Structure determination of 1,5–anhydro–D–*erythro*–hex–1–en–3–ulose (1) as a metabolite of 3”–oxo–puerarin catalyzed by DgpB–C complex

The DgpB–C complex cleaves the *C*–glycosidic bond in 3”–oxo–puerarin to produce daidzein, whereas the other metabolite corresponding to the precursor of D–glucose, derived from the sugar moiety in 3”–oxo–puerarin, remained unknown. To determine the structure of the real metabolite, enzymatic *C*–deglycosylation of 3”–oxo–puerarin was used. The major metabolite was obtained by chromatographic separation; the ^1^H and ^13^C NMR spectra are shown in Fig. 6. Based on spectral analysis, the signal for H–1 appeared at δ 7.36 ppm in the ^1^H NMR spectrum and signals for C–1, C–2, and C–3 were observed at δ 148.0, 135.2 and 191.5 ppm, respectively, in the ^13^C NMR spectrum. These results suggest that the metabolite contained an *α,β*–unsaturated carbonyl group. Further analysis and comparison of the spectral data with previously reported data (23, 24) revealed that the structure of the real metabolite was 1,5–anhydro–D–*erythro*–hex–1–en–3–ulose (**1**).

**Fig. 6.**
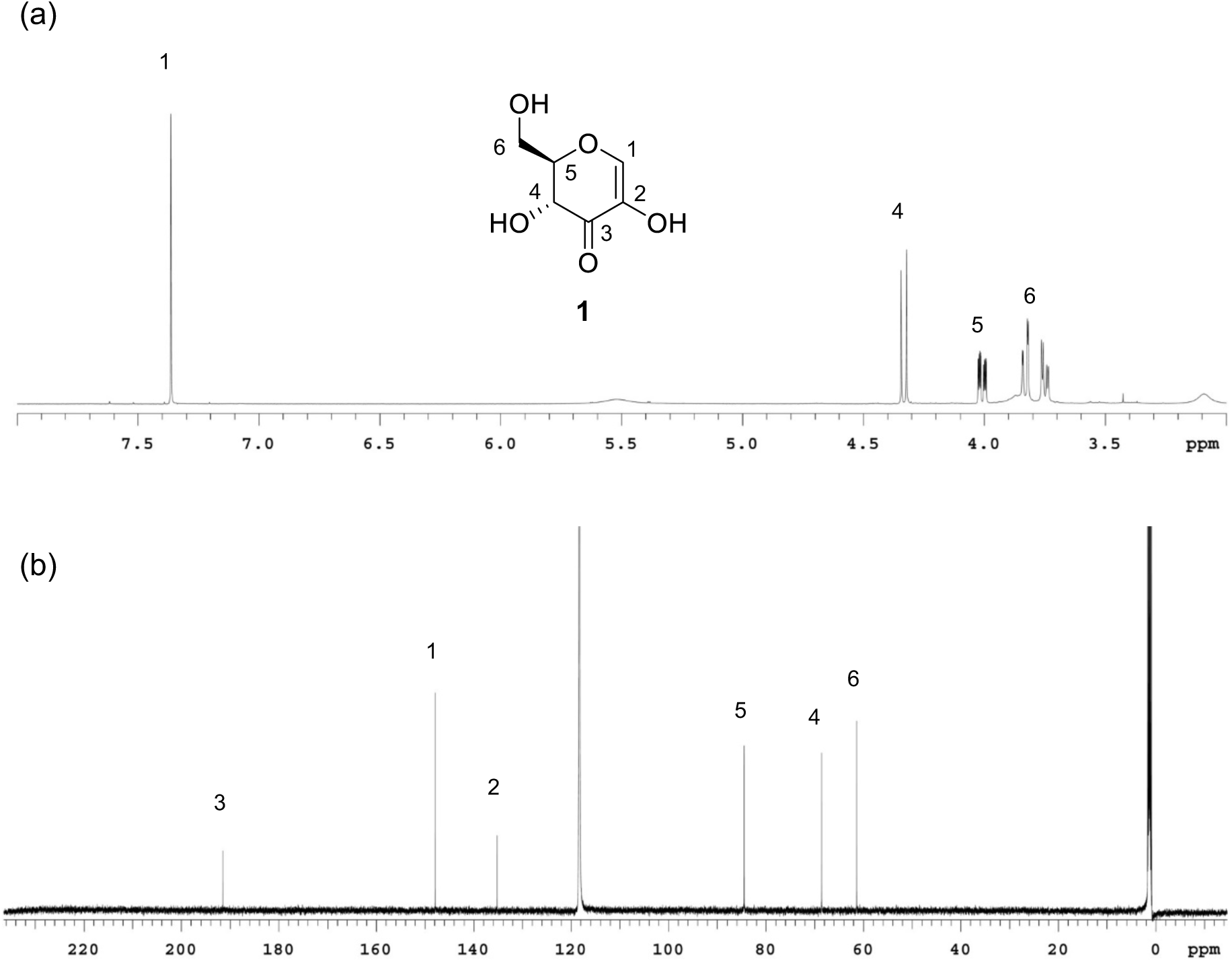
^1^H– and ^13^C–NMR spectra of **1**, a metabolite derived from 3″–oxo–puerarin sugar part catalyzed by DgpB–C complex.

## Discussion

The human intestinal bacterium strain PUE can metabolize the isoflavone *C*–glucoside puerarin to daidzein; however, the metabolic enzymes have not been well-characterized. In this study, the bacterial DgpA protein was identified as the enzyme responsible for puerarin oxidation, as shown in Fig. 1. Additionally, enzymatic *C*–deglycosylation of puerarin was accomplished by a combination of recombinant DgpA, DgpB–C complex, and 3–oxo–glucose, yielding daidzein and **1**.

We previously purified a 40 kDa protein involved in puerarin metabolism from strain PUE and sequenced the 30 N-terminal amino acids (17). The sequence was identical to that of DgpA, indicating that the purified protein, previously designated as protein C, was DgpA. DgpA protein is a member of the Gfo/Idh/MocA oxidoreductase family. These proteins typically utilize NAD^+^/NADP^+^ and are related to the redox reactions of pyranoses (22). Gfo/Idh/MocA family oxidoreductases have a two-domain structure, N–terminal NAD(P)-binding Rossmann fold domain, and C–terminal *α*/*β* domain involved in substrate binding (22). Our results revealed that DgpA was an NAD(H)-binding enzyme and used 3–oxo–glucose as the hydride accepter for puerarin oxidation. Therefore, DgpA-bound NAD(H) at the N–terminal Rossmann fold domain likely plays an important role in the redox reaction between the two substrates.

The DgpB–C complex cleaved the *C*–glycosidic bond in 3”–oxo–puerarin, affording daidzein and **1** (Fig. 1). **1** was previously reported as a spontaneous decomposition product of *β*–elimination of 3–ketocarbohydrates, such as 3–ketosucrose (25), ginsenoside oxidized compound K (26), and 3–keto–levoglucosan (23) under alkaline conditions. Similar *β*–elimination-like cleavage has been observed in glycoside hydrolase families 4 and 109 (27, 28). These protein families show a unique reaction mechanism involving NAD^+^ for glycosyl bond cleavage. The first step of the reaction is oxidation at the C–3 hydroxyl of glycosides to yield 3–keto–glycosides and NADH. Consequently, the acidified C–2 proton adjacent to the C–3 ketone is abstracted, and then a glycosidic linkage is cleaved by *β*–elimination to give *α,β*–unsaturated carbonyl intermediate such as **1**. The hydrolase reaction is completed by Michael-type 1,4-addition of H_2_O to the intermediate and subsequent reduction of C–3 ketone to hydroxyl assisted by NADH. The glycoside hydrolase family 4 protein was reported to cleave not only *O*–glycosides but also more stable *S*–glycosides (29). In contrast, cleavage of *C*–glycosides by these enzymes have not been observed.

A proposed puerarin deglycosylation pathway as shown in Fig. 1 based on the above-mentioned mechanism of glycoside hydrolases 4 and 109. The other enzyme encoded in the putative puerarin-metabolizing-operon from strain PUE was likely involved in the reaction, as more than three enzymes were reported to participate in puerarin *C*–deglycosylation (17). To clarify the enzymatic puerarin *C*–deglycosylation, further studies are needed to characterize the unidentified enzyme responsible for the enantioselective Michael addition of H_2_O to **1** to provide 3–oxo–glucose and identify the other gene products DgpD–H.

## Acknowledgement

This research was supported by JSPS KAKENHI Grant Number 18K14940.

## REFERENCES

1. Sekirov I, Russell SL, Antunes LCM, Finlay BB. 2010. Gut microbiota in health and disease. Physiol Rev 90:859–904.

2. Rajilić-Stojanović M, de vos WM. 2014. The first 1000 cultured species of the human gastrointestinal microbiota. FEMS Microbiol Rev 38:996–1047.

3. Rowland I, Gibson G, Heinken A, Scott K, Swann J, Thiele I, Tuohy K. 2018. Gut microbiota functions: metabolism of nutrients and other food components. Eur J Nutr 57:1–24.

4. Braune A, Blaut M. 2016. Bacterial species involved in the conversion of dietary flavonoids in the human gut. Gut Microbes 7:216–34.

5. Setchell KD, Brown NM, Lydeking–Olsen E. 2002. The clinical importance of the metabolite equol – a clue to the effectiveness of soy and its isoflavones. J Nutr 132:3577–3584.

6. Setchell KD, Clerici C. 2010. Equol: history, chemistry, and formation. J Nutr 140:1355S–1362S.

7. Kobashi K, Akao T. 1997. Relation of intestinal bacteria to pharmacological effects of glycosides. Bioscience Microflara 16:1–7.

8. Che QM, Akao T, Hattori M, Kobashi K, Namba T. 1991. Isolation of a human intestinal bacterium capable of transforming barbaloin to aloe-emodin anthrone. Planta Med 57:15–19.

9. Sanugul K, Akao T, Li Y, Kakiuchi N, Nakamura N, Hattori M. 2005. Isolation of a human intestinal bacterium that transforms mangiferin to norathyriol and inducibility of the enzyme that cleaves a *C*–glucosyl bond. Biol Pharm Bull 28:1672–1678.

10. Jin JS, Nishihata T, Kakiuchi N, Hattori M. 2008. Biotransformation of *C*–glucosylisoflavone puerarin to estrogenic (3*S*)–equol in co–culture of two human intestinal bacteria. Biol Pharm Bull 31:1621–1625.

11. Braune A, Blaut M. 2011. Deglycosylation of puerarin and other aromatic *C*–glucosides by a newly isolated human intestinal bacterium. Environ Microbiol 13:482–494.

12. Braune A, Blaut M. 2012. Intestinal bacterium *Eubacterium cellulosolvens* deglycosylates flavonoid *C*–and *O*–glucosides. Appl Environ Microbiol 78:8151–8153.

13. Kim M, Lee J, Han J. 2015. Deglycosylation of isoflavone *C*–glycosides by newly isolated human intestinal bacteria. J Sci Food Agric 95:1925–1931.

14. Zheng S, Geng D, Liu S, Wang Q, Liu S, Wang R. 2019. A newly isolated human intestinal bacterium strain capable of deglycosylating flavone *C*–glycosides and its functional properties. Microb Cell Fact 18:94.

15. Braune A, Engst W, Blaut M. 2016. Identification and functional expression of genes encoding flavonoid *O–* and *C*–glycosidases in intestinal bacteria. Environ Microbiol 18:2117–2129.

16. Nakamura K, Nishihata T, Jin JS, Ma CM, Komatsu K, Iwashima M, Hattori M. 2011. The *C*–glucosyl bond of puerarin was cleaved hydrolytically by a human intestinal bacterium strain PUE to yield its aglycone daidzein and an intact glucose. Chem Pharm Bull 59:23–27.

17. Nakamura K, Komatsu K, Hattori M, Iwashima M. 2013. Enzymatic cleavage of the *C*–glucosidic bond of puerarin by three proteins, Mn^2+^, and oxidized form of nicotinamide adenine dinucleotide. Biol Pharm Bull 36:635–640.

18. Nakamura K, Zhu S, Komatsu K, Hattori M, Iwashima M. 2019. Expression and characterization of the human intestinal bacterial enzyme which cleaves the *C*–glycosidic bond in 3”–oxo–puerarin. Biol Pharm Bull 42:417–423.

19. Fukuyama K, Ohrui H, Kuwahara S. 2015. Synthesis of EFdA via a diastereoselective aldol reaction of a protected 3–keto furanose. Org Lett 17:828–831.

20. Vetter ND, Langill DM, Anjum S, Boisvert-Martel J, Jagdhane RC, Omene E, Zheng H, van Straaten KE, Asiamah I, Krol ES, Sanders DA, Palmer DR. 2013. A previously unrecognized kanosamine biosynthesis pathway in *Bacillus subtilis*. J Am Chem Soc 135:5970–5973.

21. Daussmann T, Aivasidis A, Wandrey C. 1997. Purification and characterization of an alcohol:N,N-dimethyl-4-nitrosoaniline oxidoreductase from the methanogen Methanosarcina barkeri DSM 804 strain Fusaro. Eur J Biochem 248:889–896.

22. Taberman H, Parkkinen T, Rouvinen J. 2016. Structural and functional features of the NAD(P) dependent Gfo/Idh/MocA protein family oxidoreductases. Protein Sci 25:778–786.

23. Kitaoka M. 2017. Synthesis of 3–keto–levoglucosan using pyranose oxidase and its spontaneous decomposition via β–elimination. J appl glycosci 64:99–107.

24. Chung K, Waymouth RM. 2016. Selective catalytic oxidation of unprotected carbohydrates. ACS Catal 6:4653–4659.

25. Pietsch M, Walter M, Buchholz K. 1994. Regioselective synthesis of new sucrose derivatives via 3–ketosucrose. Carbohydr Res 254:183–194.

26. Kim EM, Seo JH, Baek K, Kim BG. 2015. Characterization of two-step deglycosylation via oxidation by glycoside oxidoreductase and defining their subfamily. Sci Rep 5:10877.

27. Yip VL, Varrot A, Davies GJ, Rajan SS, Yang X, Thompson J, Anderson WF, Withers SG. 2004. An unusual mechanism of glycoside hydrolysis involving redox and elimination steps by a family 4 beta–glycosidase from *Thermotoga maritima*. J Am Chem Soc 126:8354–8355.

28. Jongkees SA, Withers SG. 2014. Unusual enzymatic glycoside cleavage mechanisms. Acc Chem Res 47:226–235.

29. Yip VL, Withers SG. 2006. Family 4 glycosidases carry out efficient hydrolysis of thioglycosides by an *α, β*–elimination mechanism. Angew Chem Int Ed 45:6179–6182.

